# Chemical exploration of a highly selective scaffold with activity against intracellular *Mycobacterium tuberculosis*

**DOI:** 10.1101/2022.02.18.481120

**Authors:** Samuel Njikan, Sara Ahmed, Alyssa Manning, Divya Awasthi, Yulia Ovechkina, Sultan Chowdhury, Arielle Butts, Tanya Parish

## Abstract

We previously identified a phenylthiourea series with activity against intracellular *Mycobacterium tuberculosis* using a high throughput, high content assay. We conducted a catalog structure-activity relationship study with a collection of 35 analogs. We identified several thiourea derivatives with excellent potency against intracellular bacteria and good selectivity over eukaryotic cells. Compounds had much lower activity against extracellular bacteria which was not increased by using cholesterol as the sole carbon source. Compounds were equally active against strains with mutations in QcrB or MmpL3, thereby excluding common, promiscuous targets as the mode of action. The phenylthiourea series represents a good starting point for further exploration to develop novel anti-tubercular agents.

## Introduction

Despite decades of extensive research efforts towards diagnosis, vaccination, and treatment, tuberculosis remains one of the leading infectious causes of death globally. The World Health Organization estimates that nearly one in four people are latently infected with *M. tuberculosis* and there were 1.5 million deaths in 2020 (1). Infection with *Mycobacterium tuberculosis* generally occurs through the inhalation of aerosolized bacteria released by those with active tuberculosis. Bacteria are phagocytosed in the lower respiratory tract by alveolar macrophages (2). In most cases this results in a latent infection and the individual remains asymptomatic and non-contagious. However, in some cases, active symptomatic disease develops and reactivation can occur as a result of immune suppression or increased immune stress (3).

While there are therapeutic interventions available for the treatment of disease, they leave much to be desired. Treatment for drug-susceptible *M. tuberculosis* requires a multidrug regimen taken over a period of six months. Side effects of the individual components, drug-drug interactions, and the length of treatment dramatically impact compliance thereby reducing cure rates and contributing to the emergence of drug resistant (4). Treatment of drug resistant strains requires more complex regimens over a prolonged period further reducing the likelihood of a successful outcome (5). The need for a complex treatment regimen for tuberculosis is rooted in the complex and heterogenous nature of the disease. During infection, *M. tuberculosis* exists in the host in several micro-environments and metabolic states, which need to be targeted during treatment (6).

While target-based drug discovery efforts are experiencing a resurgence in the field of antibacterial drug discovery, and in tuberculosis research in particular, the majority of successful agents are still identified through more traditional whole cell screening approaches (7,8). An understanding of the properties that contribute to cellular penetration and uptake is still lagging in the field of bacteriology, as compared to our extensive understanding of the features when it comes to mammalian cell penetration (9). Anti-mycobacterials must also penetrate the waxy mycomembrane. To identify novel scaffolds requires venturing into previously unexplored chemical space or the development of unique screening strategies with improved sensitivity and/or different specificity.

A key feature of *M. tuberculosis* infection that contributes significantly to both pathogenesis and treatment difficulty, is its ability to survive and proliferate inside macrophages (10). In the context of treatment, this means that active compounds must access this intracellular compartment to exert their antibacterial activity. This requirement is rarely considered during the compound screening phase if compounds are examined against axenically grown bacteria. To address this limitation, we developed a high content, high throughput screening assay to determine activity of compounds against *M. tuberculosis* growing within mammalian macrophages (11). Not only does this approach facilitate the prioritization of compounds with intracellular activity but it also enables us to filter based on macrophage survival and remove cytotoxic compounds from follow-up studies. Additionally, this expands the realm of potential targets to include those that are conditionally essential under the physiologically relevant conditions experienced during macrophage infection which are overlooked when screening against bacteria grown in rich medium (12–14).

We previously screened a diversity collection of 10,000 compounds and identified the phenylthiourea (PTU) series as active against intracellular bacteria (15). Ureas and thioureas are useful in a variety of diseases (16–19), since they have good pharmacokinetic properties including solubility and permeability. Several studies have demonstrated the antimycobacterial properties of thiourea and urea (20). In their simplest form, urea, thiourea and guanidine are structurally similar, although the difference in heteroatoms imparts different physicochemical properties including basicity; urea and thiourea are more neutral functional groups whereas guanidine is a strong base. Both moieties have hydrogen-bond donors (from the nitrogens) and hydrogen-bond acceptors via the thio/carbonyl, which facilitates water solubility. Phenyl thioureas generally have good physicochemical profiles with low lipophilicity and low molecular weights, as well as facile synthetic routes. Based on these properties in this study, we have further investigated this series to determine its potential for development as a novel anti-tubercular agent.

## Results

### Series identification and expansion

We previously demonstrated that PTU analogs have some potential as anti-tubercular agent since our previous work demonstrated that 4/5 analogs in the screening collection had activity (Table 1) (15). Three of these compounds (**1, 2**, and **5**) had excellent potency and selectivity for *M. tuberculosis* over eukaryotic cells (Table 1) (15). The thiourea analogs demonstrated activity against extracellular bacteria but were 5-10-fold less active (15). Compounds **1** and **2** were thioureas with excellent intracellular activity (IC_50_ of 0.4 and 0.3 μM respectively) (15). Since the series targeted both intracellular and extracellular bacteria, we considered this to be a good starting point for exploration.

**Table 1.**
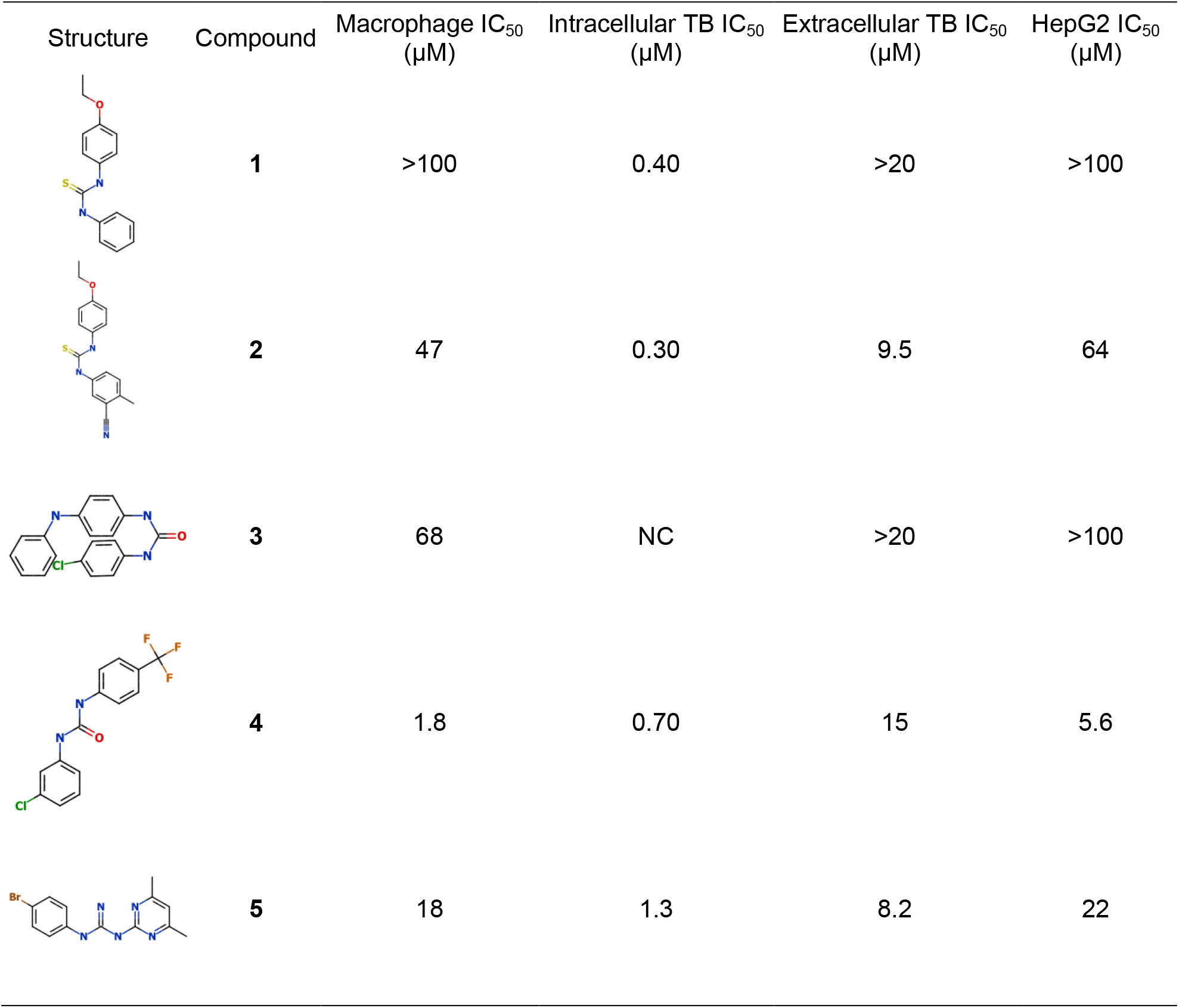
Phenylthiourea hits identified from the primary screen. *Activity against macrophages (RAW 264.7) and intracellular *M. tuberculosis* were determined using the high content assay. ^#^Activity against extracellular *M. tuberculosis* or HepG2 cells was determined in standard culture medium. IC_50_ = concentration required to inhibit by 50%. Data are the average of at least two independent experiments. NC - not calculated due to cytotoxicity. Taken from (15).

| Structure | Compound | Macrophage IC<sub>50</sub><br>(μM) | Intracellular TB IC<sub>50</sub><br>(μM) | Extracellular TB IC<sub>50</sub><br>(μM) | HepG2 IC<sub>50</sub><br>(μM) |
| --- | --- | --- | --- | --- | --- |
|  | **1** | >100 | 0.40 | >20 | >100 |
|  | **2** | 47 | 0.30 | 9.5 | 64 |
|  | **3** | 68 | NC | >20 | >100 |
|  | **4** | 1.8 | 0.70 | 15 | 5.6 |
|  | **5** | 18 | 1.3 | 8.2 | 22 |

### Characterization and structure activity relationships

We initiated a structure-activity relationship study using commercially available analogs. We identified 35 additional analogs which represented molecules containing either a thiourea core, as present in the initial hits, or a urea core. Modifications on both sides of the central core were explored including decoration of the phenyl rings, substitutions with other cyclic structures (aromatic and non-aromatic), as well as removal of one side (Table 2).

**Table 2.**
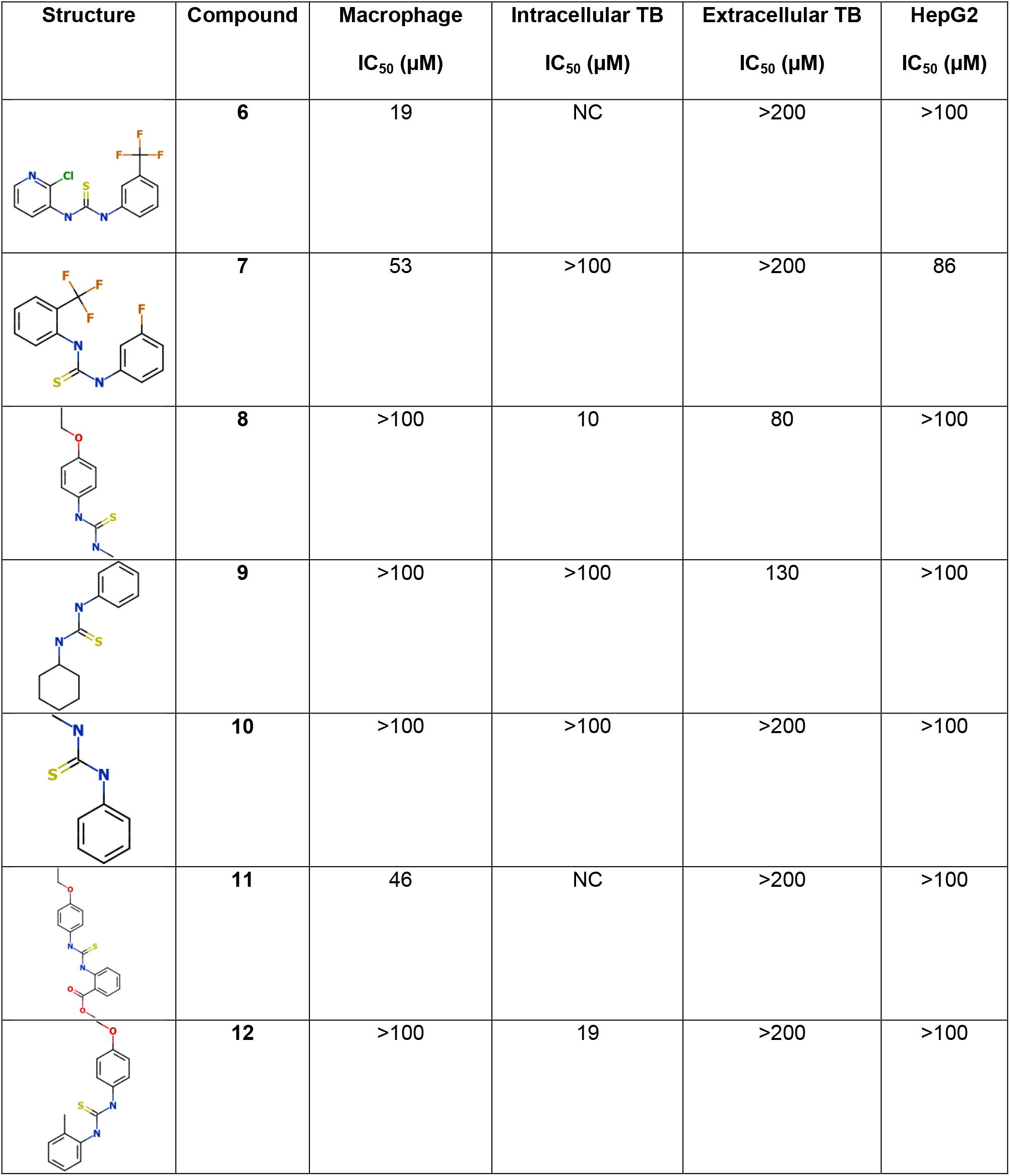
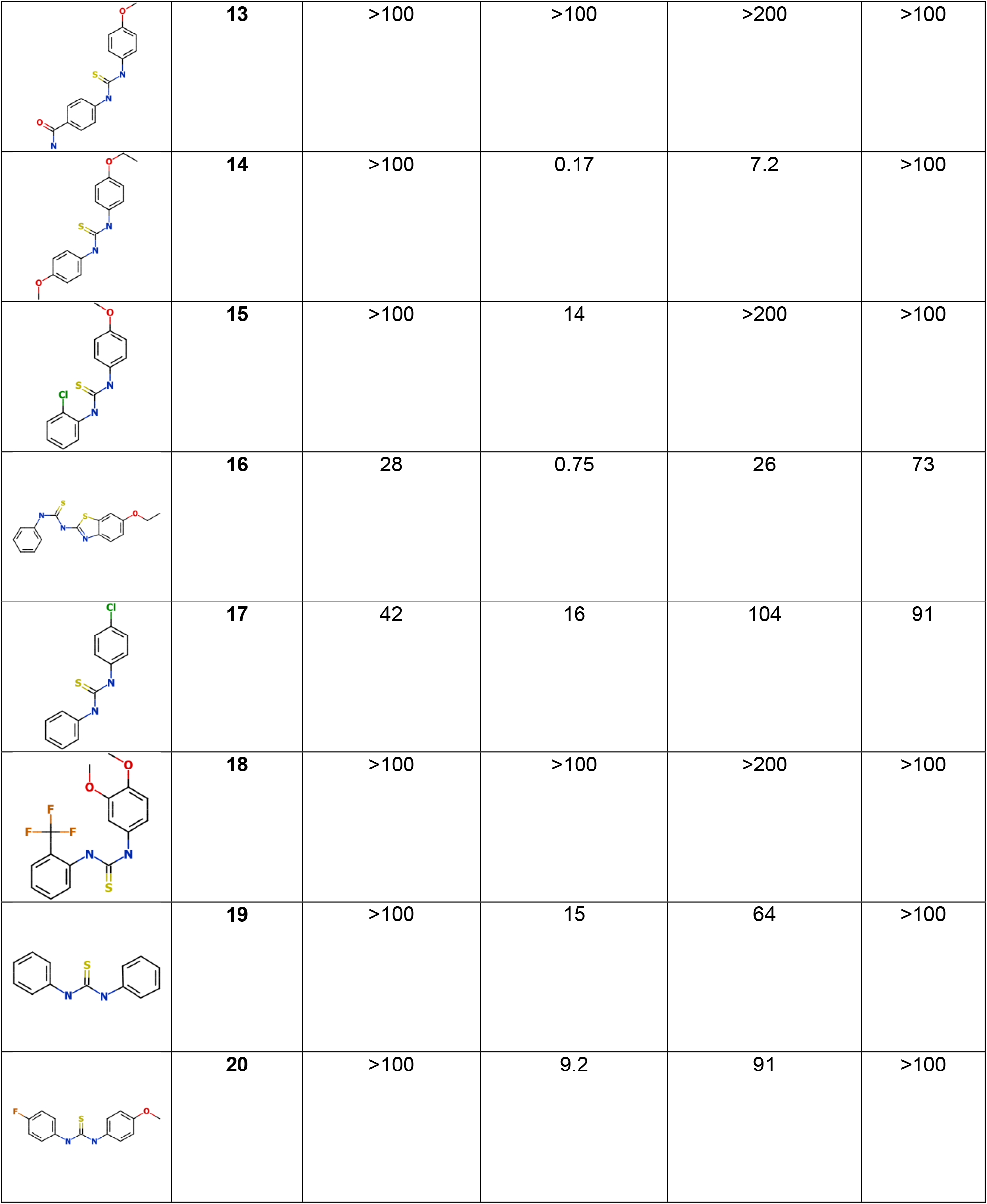
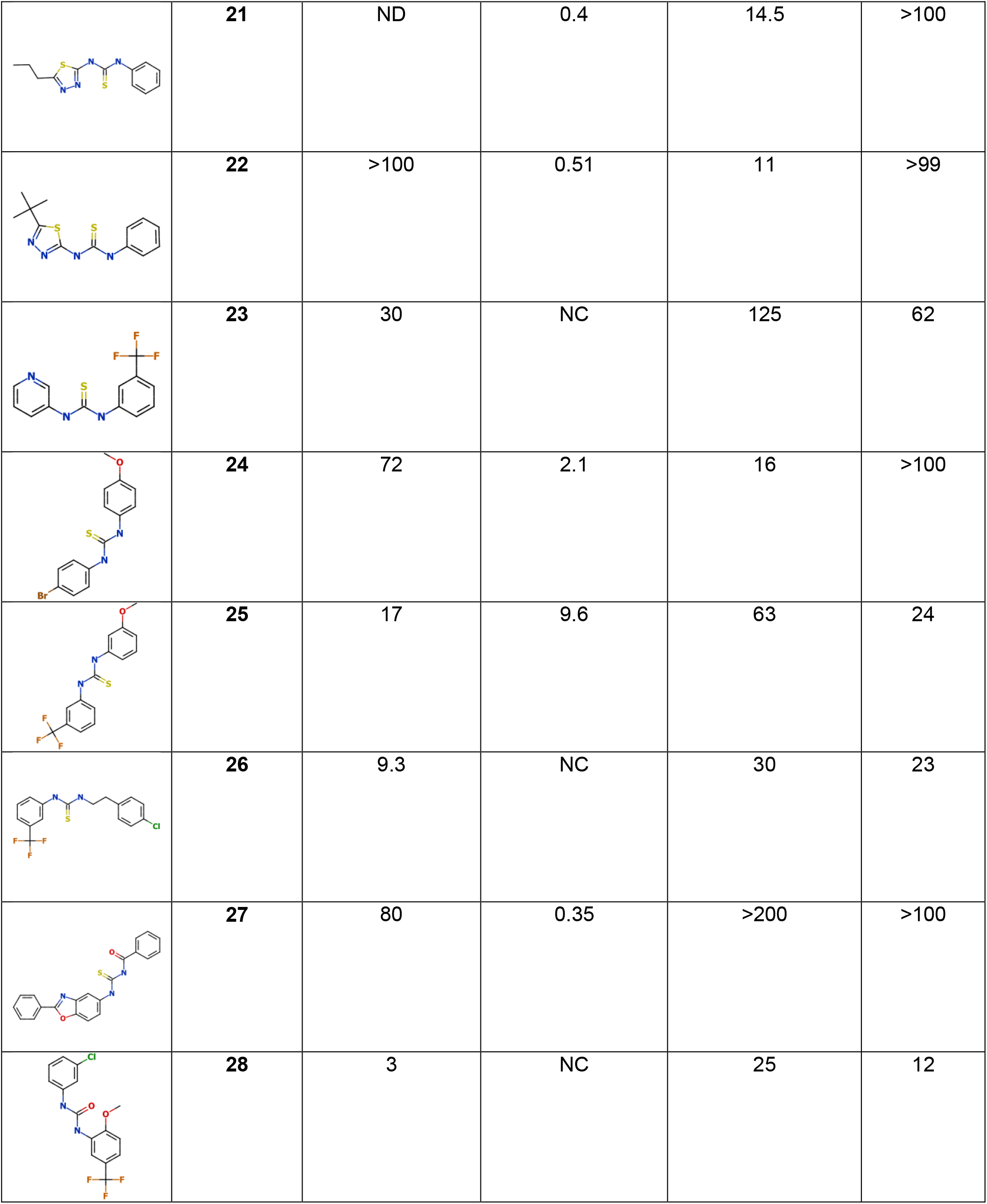
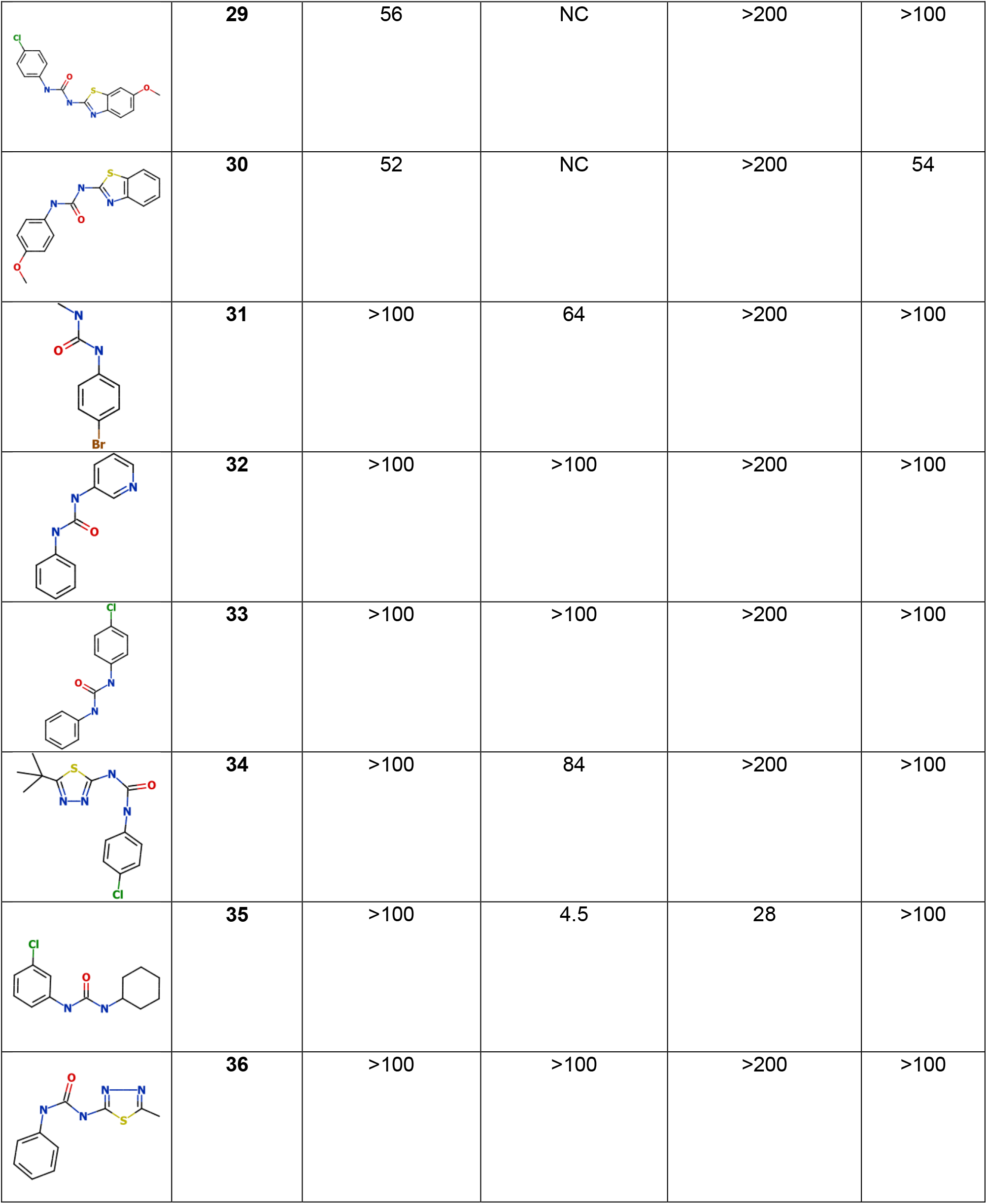
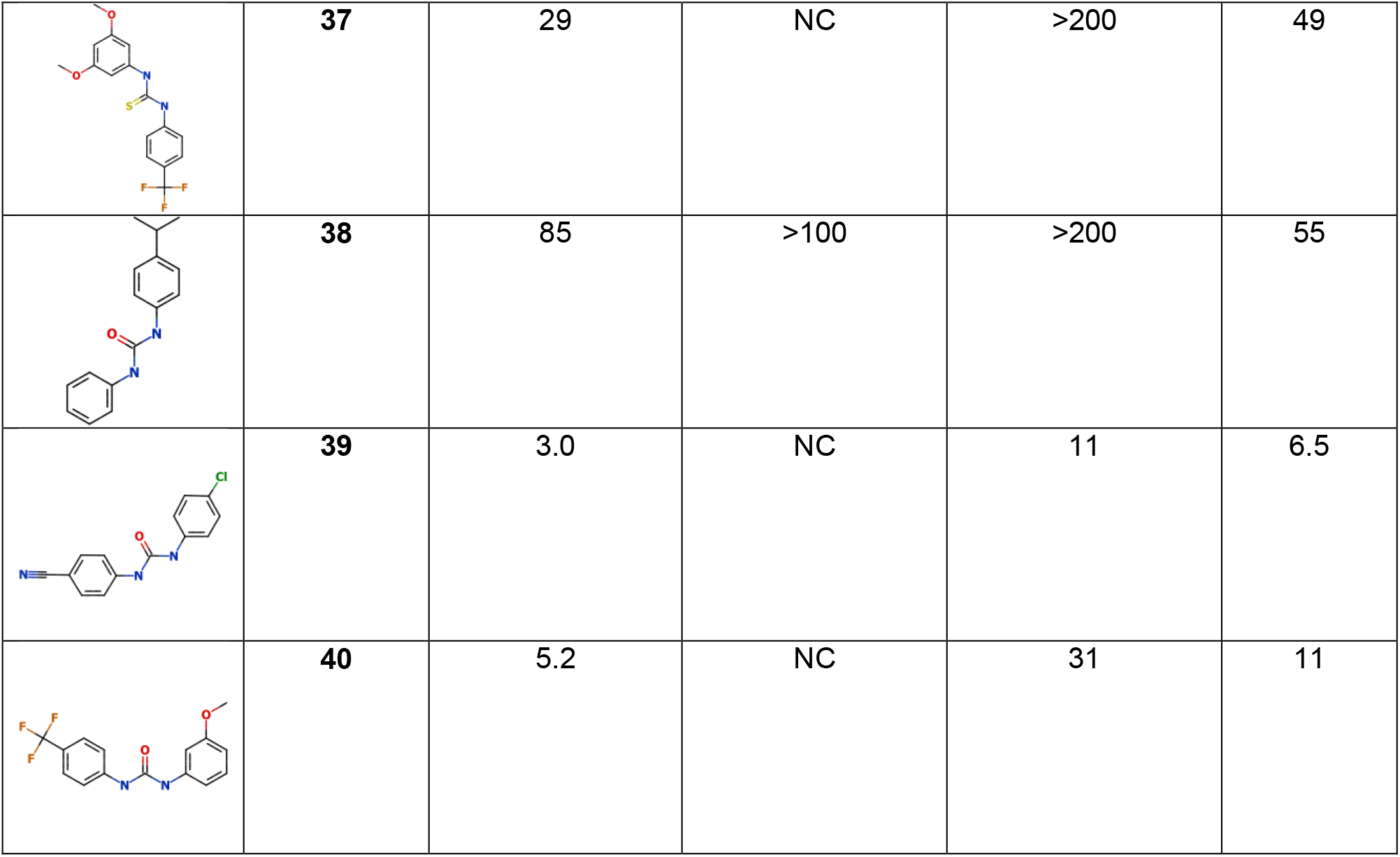
Biological activity of urea and thiourea analogs. The thiourea containing one phenyl ring with an ethoxy group at the C-4 position of the phenyl ring and an alkyl group (compound **8**) was active (IC_50_ = 10 μM) but losing the ethoxy group on the phenyl ring resulted in loss of activity (compound **10**, IC_50_ > 100 μM). A similar trend was observed with compound **9**, where the absence of the ethoxy moiety on the phenyl ring and a cyclohexyl group present on the thiourea core led to loss of intracellular activity. Ureas containing a substituted phenyl ring and an alkyl chain retained activity (compound **31**, IC_50_ = 64 μM and **35**, IC_50_ = 4.5 μM). Compound **35** had good biological activity, with some extracellular activity (IC_50_ = 28 μM) and no cytotoxicity. *Activity against macrophages (RAW 264.7) and intracellular *M. tuberculosis* was determined using the high content assay. ^#^Activity against extracellular *M. tuberculosis* or HepG2 cells was determined in standard culture medium. IC_50_ = concentration required to inhibit by 50%. Data are the average of at least two independent experiments. NC - not calculated due to cytotoxicity.

| Structure | Compound | Macrophage<br>$\text{IC}_{50} (\mu\text{M})$ | Intracellular TB<br>$\text{IC}_{50} (\mu\text{M})$ | Extracellular TB<br>$\text{IC}_{50} (\mu\text{M})$ | HepG2<br>$\text{IC}_{50} (\mu\text{M})$ |
| --- | --- | --- | --- | --- | --- |
|  | **6** | 19 | NC | >200 | >100 |
|  | **7** | 53 | >100 | >200 | 86 |
|  | **8** | >100 | 10 | 80 | >100 |
|  | **9** | >100 | >100 | 130 | >100 |
|  | **10** | >100 | >100 | >200 | >100 |
|  | **11** | 46 | NC | >200 | >100 |
|  | **12** | >100 | 19 | >200 | >100 |
|  | **13** | >100 | >100 | >200 | >100 |
|  | **14** | >100 | 0.17 | 7.2 | >100 |
|  | **15** | >100 | 14 | >200 | >100 |
|  | **16** | 28 | 0.75 | 26 | 73 |
|  | **17** | 42 | 16 | 104 | 91 |
|  | **18** | >100 | >100 | >200 | >100 |
|  | **19** | >100 | 15 | 64 | >100 |
|  | **20** | >100 | 9.2 | 91 | >100 |
|  | **21** | ND | 0.4 | 14.5 | >100 |
|  | **22** | >100 | 0.51 | 11 | >99 |
|  | **23** | 30 | NC | 125 | 62 |
|  | **24** | 72 | 2.1 | 16 | >100 |
|  | **25** | 17 | 9.6 | 63 | 24 |
|  | **26** | 9.3 | NC | 30 | 23 |
|  | **27** | 80 | 0.35 | >200 | >100 |
|  | **28** | 3 | NC | 25 | 12 |
|  | **29** | 56 | NC | >200 | >100 |
|  | **30** | 52 | NC | >200 | 54 |
|  | **31** | >100 | 64 | >200 | >100 |
|  | **32** | >100 | >100 | >200 | >100 |
|  | **33** | >100 | >100 | >200 | >100 |
|  | **34** | >100 | 84 | >200 | >100 |
|  | **35** | >100 | 4.5 | 28 | >100 |
|  | **36** | >100 | >100 | >200 | >100 |
|  | **37** | 29 | NC | >200 | 49 |
|  | **38** | 85 | >100 | >200 | 55 |
|  | **39** | 3.0 | NC | 11 | 6.5 |
|  | **40** | 5.2 | NC | 31 | 11 |

We tested all 35 analogs as a ten-point, dose response curve using the high content infection assay (Table 2 and Figure 1). We determined the IC_50_ (defined as the concentration effecting 50% growth inhibition) against intracellular *M. tuberculosis* and the infected macrophages in the same samples. Compounds were also tested in ten-point dose response for activity against axenically-grown *M. tuberculosis* and cytotoxicity against the human HepG2 hepatic cell line. Compounds had a good dynamic range of activity from inactive (>100 μM for 9 molecules) to sub-micromolar (4 molecules) (Figure 1A). The most potent molecule was **14** with an IC_50_ of 0.17 μM against intracellular *M. tuberculosis*. Notably the four most active molecules were all thioureas (Figure 1A). The activity of 10 molecules, representing both urea and thiourea cores, could not be determined due to cytotoxicity.

**Figure 1.**
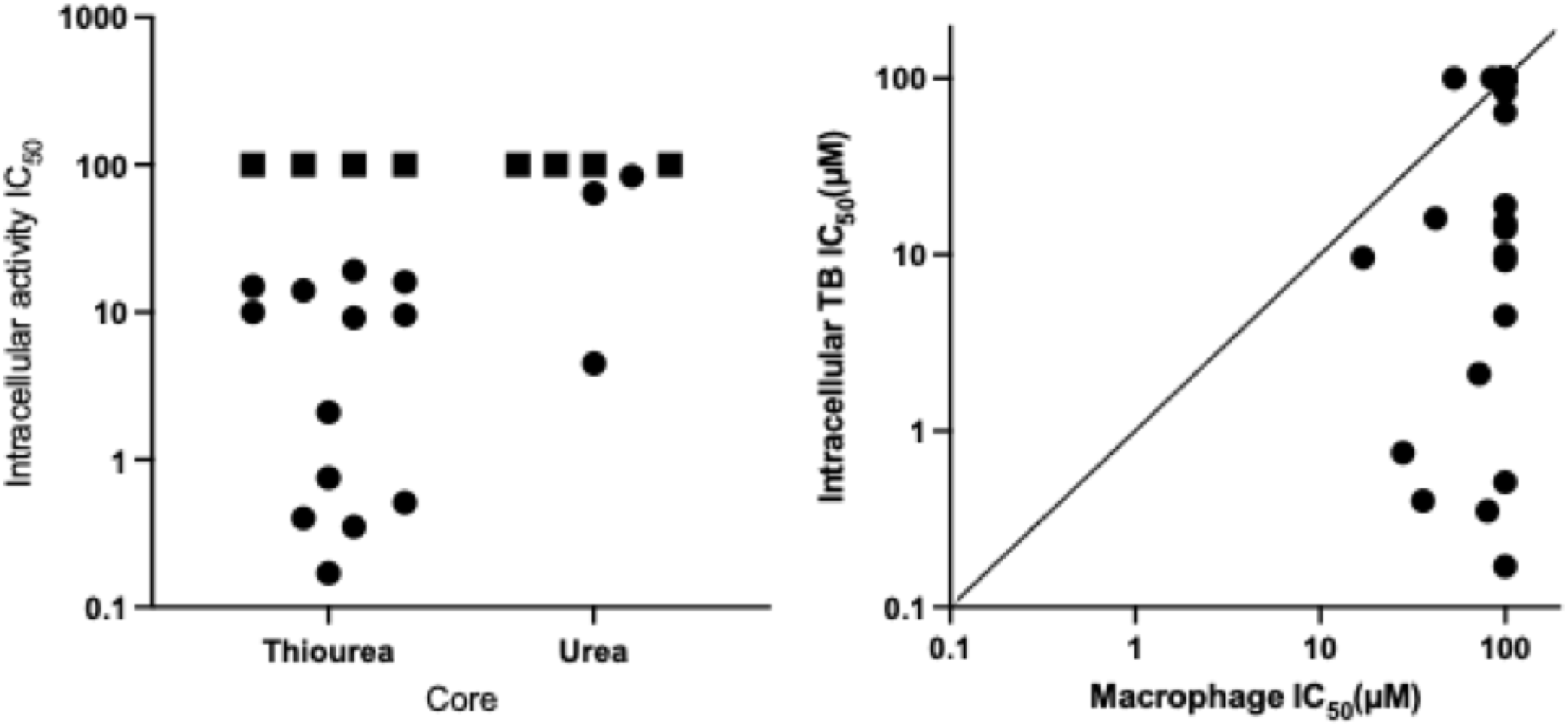
Anti-tubercular activity of ureas and thioureas against intracellular *M. tuberculosis*. (A) Anti-tubercular activity by core. Squares indicate where no activity was seen i.e IC_50_ > μM. (B) Activity against *M. tuberculosis* and murine macrophages. Activity against macropages (RAW 264.7) and intracellular *M. tuberculosis* was determined using the high content assay. IC_50_ = concentration required to inhibit by 50%. Data are the average of at least two independent experiments.

Analogs **8, 12, 15, 17** and **19** containing a thiourea pharmacophore exhibited good potency (IC_50_ <20 μM) and five thiourea compounds (**14, 16, 21, 22** and **27**) had excellent potency (IC_50_ <1.0 μM) (Table 2). The common feature in these asymmetrical thiourea analogs is the presence of an electron-rich phenyl ring attached to either side of the thiourea functionality. Interestingly, while several of the analogs were also active against extracellular bacteria, the potency was greatly reduced; for example, compound **14** was the most active with an IC_50_ of 7.2 μM against extracellular bacteria, and IC_50_ of 0.17 μM against intracellular bacteria (>40-fold difference).

Compounds with varied combinations of electron-withdrawing groups and electron-donating group at either side of the thiourea moiety showed very different biological activity. Compound **15** with a methoxy group attached to one phenyl ring and a chlorine atom on the other phenyl ring and compound **17** (IC_50_ = 16 μM) which lacked the methoxy were equally active (IC_50_ of 14 and 16 μM respectively). Compound **13** with amide and methoxy groups was inactive, as was compound **18** with a trifluoromethyl group and two electron-rich methoxy groups. However, similar analogs with different combinations were active; analogs **20** and **25** with a strong electron-withdrawing group (fluorine or trifluoromethyl) on one phenyl ring and an electron rich methoxy group on the other phenyl ring had activity (IC_50_ = 9.2 μM and IC50 = 9.6 μM respectively). It is worth mentioning in this perspective that the attachment of the trifluoromethyl group was different in each compound. However, replacement of the strong electron withdrawing fluorine atom with a weaker electron-withdrawing group (bromine) (compound **24**) improved activity substantially (five-fold increase in potency, IC_50_ = 2.1 μM). A phenyl ring with electron-withdrawing groups on either end of the thiourea functionality resulted in cytotoxicity to macrophages (compounds **6** and **23**). Taken together, these data suggest that compounds with an electron-rich group attached at the 4-position of the phenyl are more potent than substituents attached on either on 3- or 2-positions.

Another interesting feature in this series is seen with compounds **21, 22** and **34**; these are all asymmetrical analogs containing a 1,3,4-thiadiazole with a lipophilic chain attached at the C-5 position. Compounds **21** and **22** with the thiourea functionality are potent (IC_50_ = 0.40 μM and IC_50_ = 0.51 μM respectively), whereas compound **34** with a urea group lost activity. The presence of an unsubstituted phenyl ring and a heteroaromatic 1,3,4-thiadiazole ring at one end of the thiourea functional group provides a future opportunity to explore substitutions in different positions as well as to expand to other 5-member heteroaromatic ring systems in order to improve drug-like properties of the molecule. A direct comparison between compound **22** (IC_50_ = 0.51 μM) and compound **34** (IC_50_ = 84 μM) reveals the requirement for the thiourea moiety.

Compound **27** (IC_50_ = 0.35 μM) has a unique acyl thiourea functional group. The asymmetrical acyl thiourea provides a handle to explore both sides of the molecule by addition of different functional moieties to improve lipophilicity and molecular weight. This opens possibilities to prepare derivatives of compound **27** in order to explore different 5- and 6-member rings to improve physicochemical properties.

### Cytotoxicity against HepG2 correlates with cytotoxicity against RAW 264.7 cells

There was a strong correlation between cytotoxicity against HepG2 cells and infected macrophages (Figure 2A). Several compounds showed a combination of potency against intracellular *M. tuberculosis* and good selectivity (Table 2 and Figure 1B). For example, compound **14** has sub-micromolar activity (IC_50_ = 0.17 μM) with a lack of cytotoxicity against either cell line (IC_50_ >100 μM) giving a selectivity index (SI) of 588. Similarly, compounds **21, 22** and **27** all had SI of > 200. Seventeen compounds showed no activity against either eukaryotic cell line (Table 2). In addition, activity against *M. tuberculosis* was not correlated with cytotoxicity, demonstrating that the series does not have a cytotoxicity issue.

**Figure 2.**
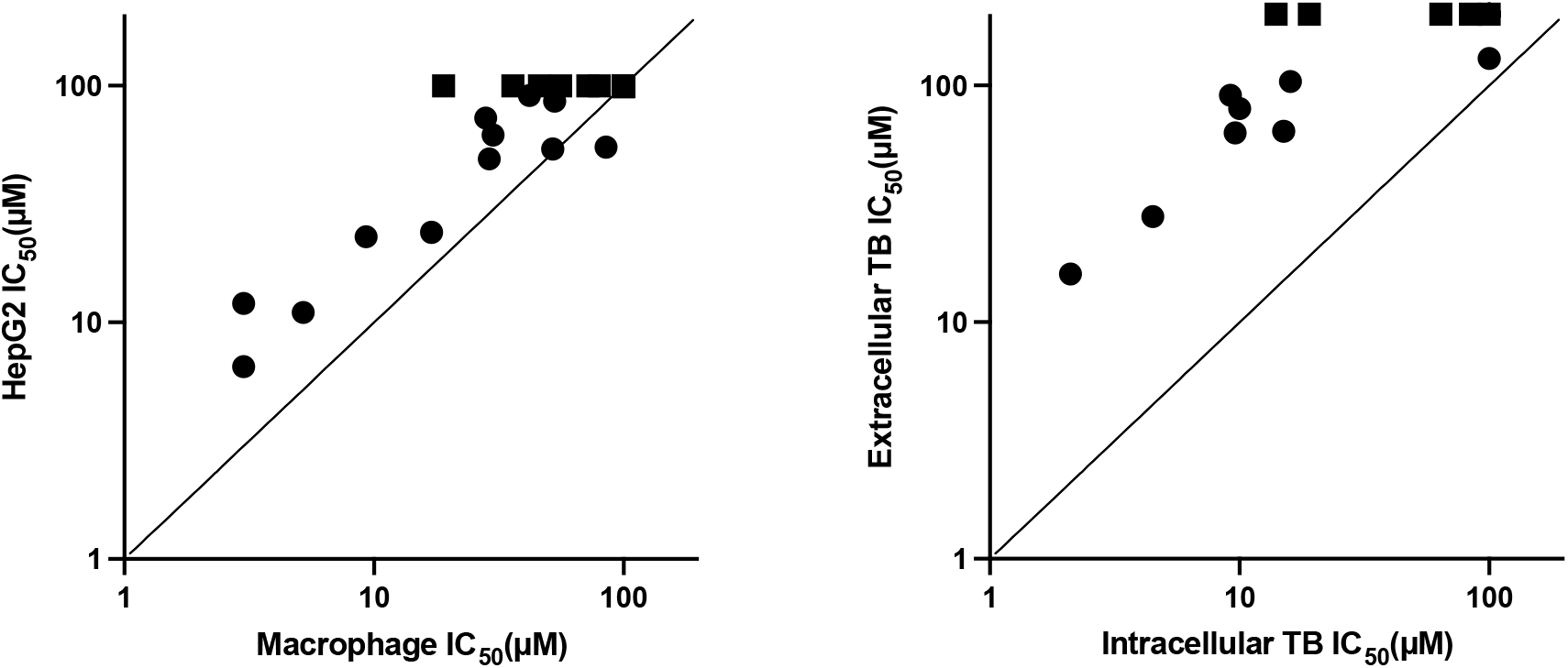
Biological profile of PTU series. (A) Comparison of cytotoxicity against RAW267.4 and HepG2 cells. (B) Comparison of activity against intracellular and extracellular *M. tuberculosis*. Activity against macrophages (RAW 264.7) and intracellular *M. tuberculosis* was determined using the high content assay. Activity against extracellular *M. tuberculosis* or HepG2 cells was determined in standard culture medium. IC_50_ = concentration required to inhibit by 50%. Data are the average of at least two independent experiments. Squares indicate where no activity was seen i.e IC_50_ >100 μM (>200 μM for the extracellular bacterial assay)

### PTU compounds are more active against intracellular bacteria

We tested all analogs for activity against axenically cultured *M. tuberculosis*. (Table 2 and Figure 2B). Several of the analogs with intracellular activity were also active against extracellular bacteria, but with much lower potency, ranging from a 4.3-fold difference (**19** with intracellular IC_50_ of 15 μM and extracellular IC_50_ of 64 μM) to more than 20-fold (**22** with intracellular IC_50_ of 0.51 μM and extracellular IC_50_ of 11 μM). Compound **27** had the largest differential activity of >500-fold with intracellular IC_50_ of 0.35 μM and no extracellular activity (IC_50_ >200 μM).

### PTUs exhibit minimal activity against extracellular bacteria

To further assess the activity of this series, three analogs with a broad range of potencies were selected for kill kinetics under standard replicating conditions (Figure 3). Compounds were tested up to 100 μM with minimal impact on culture viability over 21 days. Even at the highest concentrations, there was no inhibition of growth in the first 7 days consistent with the results seen in the 5-day assay. One compound (**14)** resulted in a small decrease in viable bacteria at day 21 at the highest concentration. Thus, we confirmed that compounds have minimal activity against axenically-cultured bacteria.

**Figure 3.**
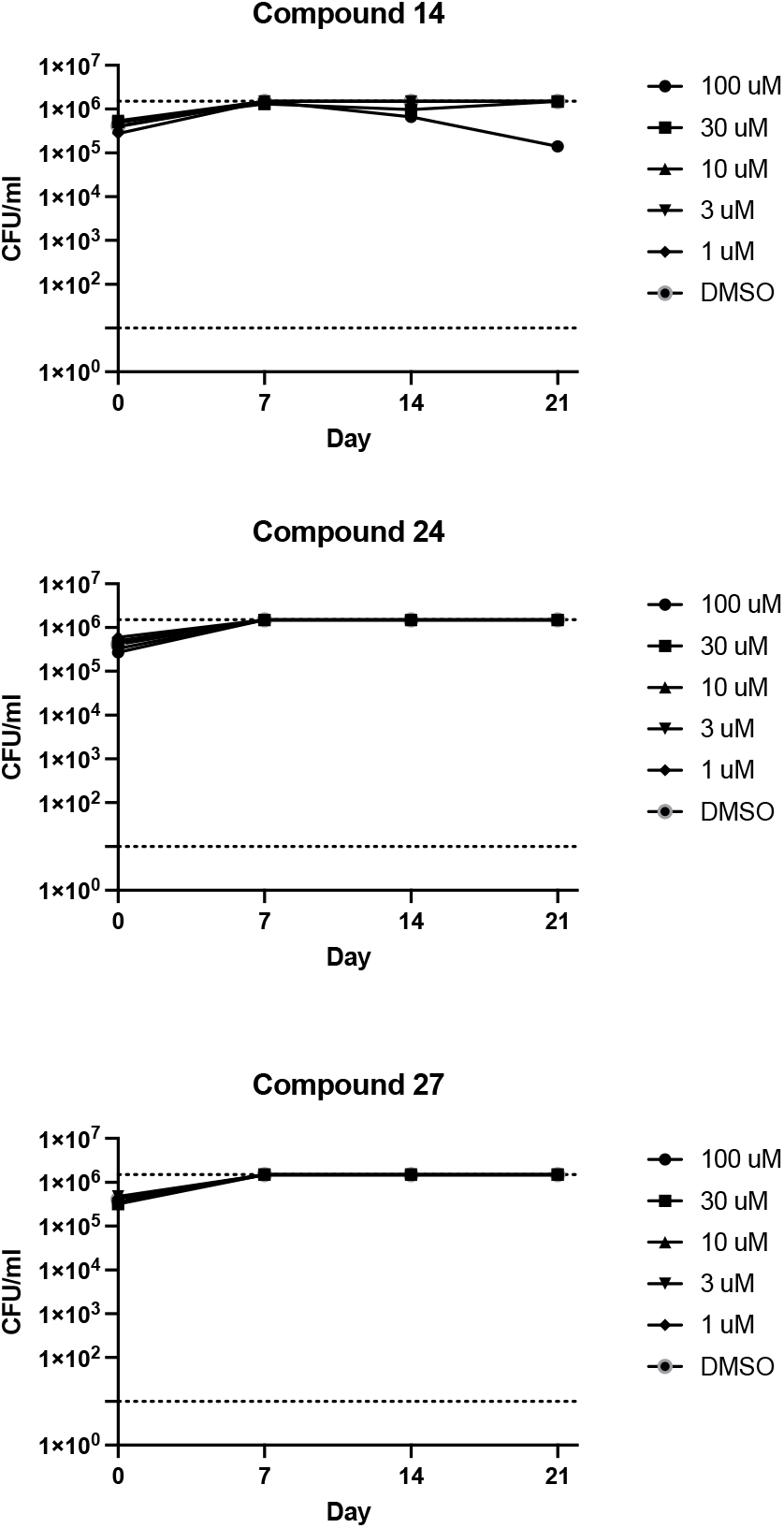
Kill kinetics of representative compounds against replicating *M. tuberculosis*. *M. tuberculosis* was exposed to compounds under replicating conditions (aerobic culture) in 7H9-Tw-OADC. Viability was monitored by determining CFUs.

### Target and mechanism of action

Due to the promiscuous nature of several drug targets in *M. tuberculosis* (21), we tested three selected PTU compounds against a set of strains with mutations in the most promiscuous targets. We generated fluorescent versions (*Ds*Red) of well characterized strains carrying mutations in either QcrB or MmpL3. For QcrB, we used strains with either M342T, T313I, or A396T mutations (22); for MmpL3 we used strains with a triple mutation of F255L, V646M, F644I (23). We determined IC_50_s for a small set of compounds with good activity against these strains (Figure 4). No shift in susceptibility was observed with any of the mutants tested indicating that the PTU series does not exert its antibacterial activity through these established targets.

**Figure 4.**
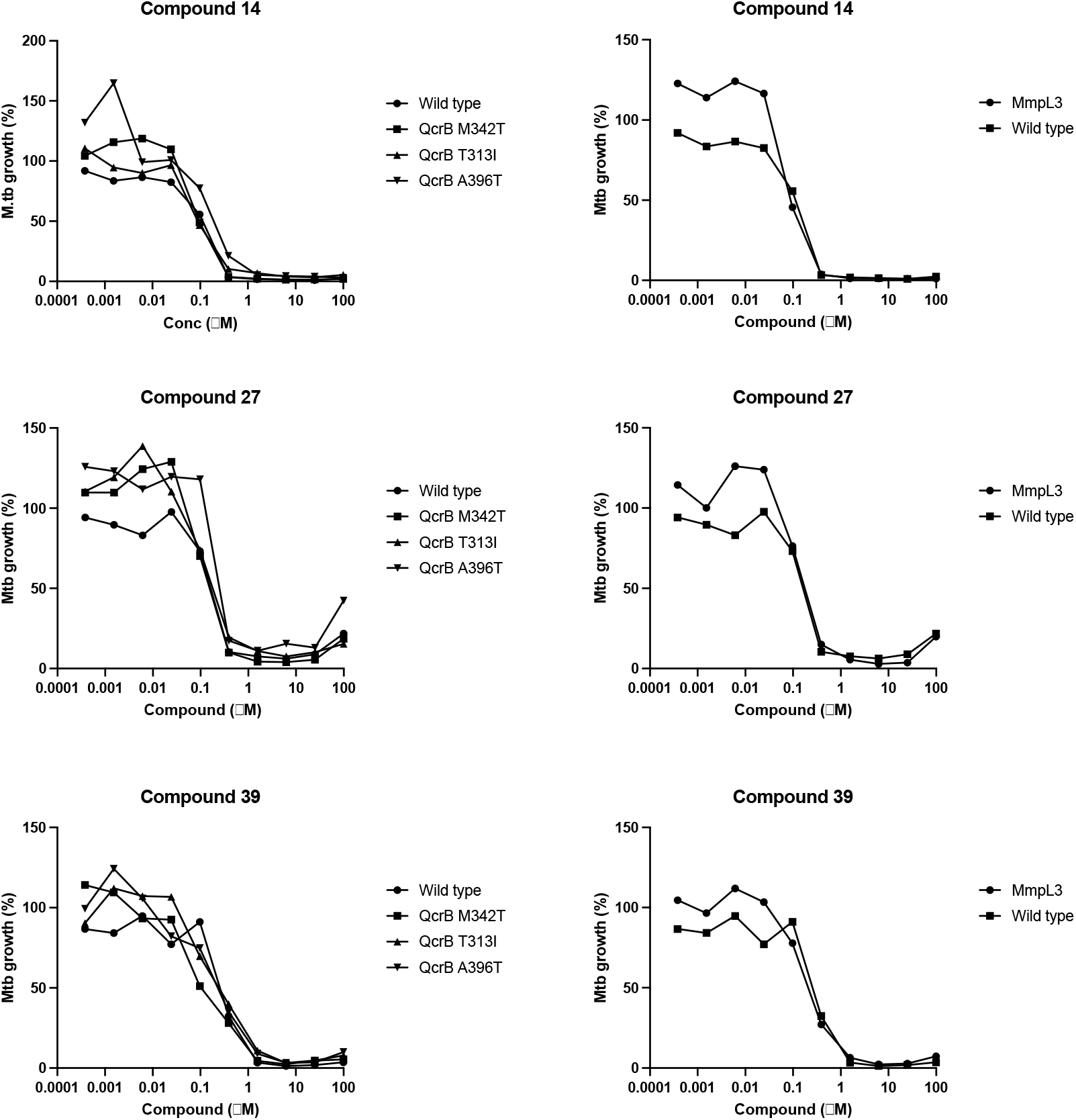
Activity against mutant strains of *M. tuberculosis*. Activity against intracellular *M. tuberculosis* strains carrying the indicated mutationswas determined using the high content assay with.

One of the best characterized, conditionally essential pathways for *M. tuberculosis* intracellular growth is cholesterol metabolism (24,25). The ability to utilize cholesterol is crucial for *M. tuberculosis* during macrophage infection (24–26). We determined whether the PTU compounds had increased activity in medium with cholesterol as the only carbon source. We tested three active compounds (Table 3). Two compounds showed no significant difference in activity from bacteria cultured with glucose (less than two-fold change either way). One compound showed marginal increase in activity (3.9-fold more active), but this was slightly below our cut-off of 4-fold being a significant difference. These small changes indicate that inhibition of cholesterol metabolism is not a mechanism of action of this series.

**Table 3.** Biological activity against extracellular *M. tuberculosis* in cholesterol medium. *Activity against extracellular *M. tuberculosis* was determined in culture medium containing either cholesterol or glucose as a carbon source. IC_50_ = concentration required to inhibit by 50%. Data are the average of at least two independent experiments.

| Compound | Cholesterol*<br>IC<sub>50</sub> (μM) | Glucose*<br>IC<sub>50</sub> (μM) | Fold-change |
| --- | --- | --- | --- |
| 14 | 15 | 7.2 | 0.5 |
| 16 | 6.6 | 26 | 3.9 |
| 22 | 5.7 | 11 | 1.9 |

## Discussion

We explored the potential of the PTU series as anti-tubercular agents targeting intracellular bacteria. We tested 35 analogs which were chosen to determine key features required for activity and selectivity. We explored the role of the central thiourea and the extent to which the sides groups could be replaced, substituted, and removed. We found that the thiourea could be replaced with a urea and maintain activity but that this was detrimental to selectivity. We determined that the two phenyl groups are not both required for activity, but only replacements with aromatic groups were tolerated. Decoration of the phenyl rings with electron withdrawing groups had a positive impact on activity with minimal effect on selectivity. Future exploration of this series would include plans to maintain the thiourea core, expand the number of aromatic groups, and focus efforts on optimizing properties with additional substitutions.

The PTU compounds exhibit activity against extracellular bacteria although to a lesser extent. This dual activity makes them attractive and is likely related to the mechanism of action of this series. Our initial mode of action investigation was designed to rule out common or well-established mechanisms. We were able to exclude common targets which suggests that the PTU compounds exert antitubercular activity through a distinct or novel target and/or mechanism. Future work will focus on target identification and exploring the range of conditions under which this series is active.

## Methods

### Culture

Murine RAW 264.7 macrophages (ATCC TIB-71) were cultured in RPMI 1640 medium supplemented with 1 mM sodium pyruvate, 2mM GlutaGro (Corning), and 5% fetal bovine serum. Human hepatic cells HepG2 (ATCC HB-8065) were cultured in DMEM with 10% fetal bovine serum and 1x penicillin streptomycin solution (100 U/ml). Cultures were maintained in a humidified incubator with 5% CO_2_ at 37°C. *M. tuberculosis* strains constitutively expressing codon-optimized DsRed from plasmid pBlazeC8 (DREAM8) (27) were cultured at 37 °C in Middlebrook 7H9 medium containing 10% v/v OADC (oleic acid, dextrose, catalase) supplement (Becton Dickinson) and 0.05% w/v Tween 80 (7H9-Tw-OADC) plus 100 μg/mL hygromycin B. *M. tuberculosis* H37Rv-LP (ATCC 25618) was the wild type strain; strains containing mutations in QcrB (M342T, T313I, or A396T) (22) or MmpL3 (23) were derived from this parental strain as described. For growth on cholesterol, *M. tuberculosis* was cultured in Middlebrook 7H9 medium containing 0.1% cas-amino acids, 500 mg/L 2-morpholinoethanesulfonic acid, 5 g/L BSA, 0.05% w/v Tyloxapol, 0.1 mM cholesterol (7H9-Tyl-Chol). Viable bacteria were determined by serial dilution and counting CFUs on Middlebrook 7H10 agar containing 10% v/v OADC after 21-28 days.

### Determination of intracellular activity

Activity against *M. tuberculosis* and RAW 264.7 cells was determined as described (11). Briefly, assay plates were prepared in clear bottom, black 384-well plates with 30 μL cRPMI and 0.6 μL compound (final concentration 1% DMSO); compounds were tested as 10-point serial dilution to generate dose response curves. 1% DMSO was used as the negative control, 10 mM isoniazid (INH) and 100 μM staurosporine (STA) were included as positive controls (maximum inhibition) for anti-tubercular activity and cytotoxicity respectively. INH and STA were also included in dose response on each plate. RAW 264.7 cells were infected with *M. tuberculosis* DREAM8 at an MOI of 1 for 24 h and extracellular bacteria removed by washing. Cells were recovered using Accumax, harvested, washed and resuspended in serum-free RPMI; 30 μL of infected cells were dispensed into each well at 3300 cells/well. Plates were incubated for 72 hours, 10 μL of 5X SYBR Green I was added, and plates were imaged with an ImageXpress Micro High Content Screening System (Molecular Devices) using a 4x objective and FITC and Texas Red channels. MetaXpress was used to analyze images. The integrated intensity of *M. tuberculosis* or macrophages was calculated for each well. Growth inhibition was calculated for each test well by normalizing to the average integrated intensity of the DMSO control wells. Curves were fitted using the Levenberg–Marquardt algorithm and IC_50_ was calculated as the compound concentration required to reduce bacterial or eukaryotic cell growth by 50%.

### Cytotoxicity

Cytotoxicity against HepG2 cells was measured after 72 hours. Cells were seeded in 384-well plates at 1800 cells per well. Compounds were added as a 10-point three-fold serial dilution after 24 h (final assay concentration of 1% DMSO). CellTiter-Glo® reagent (Promega) was added and relative luminescence units (RLU) measured. Data were normalized to the DMSO controls. Curves were fitted using the Levenberg–Marquardt algorithm; IC_50_ was calculated as the compound concentration required to reduce HepG2 cell s by 50%.

### Minimum inhibitory concentration

Inhibitory concentrations (IC_50_) against *M. tuberculosis* were determined in liquid medium as described (28). Bacterial growth was measured after 5 days by OD_590_. IC_50_ was defined as the concentration of compound required to inhibit growth *of M. tuberculosis* by 50% and was determined using the Levenberg–Marquardt least-squares plot. For MICs in cholesterol, *M. tuberculosis* was pre-cultured in 7H9-Tyl-Chol for 7 days and MICs determined after 7 days growth in 7H9-Tyl-Chol.

## Acknowledgements

We thank Lindsay Flint, Jennifer Geist, Megha Gupta, Junitta Guzman, Douglas Joerss, James Johnson, Sarah Heidl, Aaron Korkegian, Anuradha Kumar, Allison Morley, Michael Ortego, David Roberts, Bjorn Sunde, Dean Thompson, Anisa Tracy, and James Vela for technical assistance. We thank Alisha Contractor for helpful discussion.

